# Set-size effects in change detection depend on failures of retrieval and/or comparison and not on perception, encoding or storage

**DOI:** 10.1101/2020.01.19.911867

**Authors:** James C. Moreland, John Palmer, Geoffrey M. Boynton

## Abstract

Set-size effects in change detection is often used to investigate the capacity limits of dividing attention. Such capacity limits have been attributed to a variety of processes including perception, memory encoding, memory storage, memory retrieval, comparison and decision. In this study, we investigated the locus of the effect of increasing set size from 1 to 2. To measure purely attentional effects and not other phenomena such as crowding, a precue was used to manipulate relevant set size and keep the display constant across conditions. The task was to detect a change in the orientation of 1 or 2 Gabor patterns. The locus of the capacity limits was determined by varying when observers were cued to the only stimulus that was relevant. We began by measuring the baseline set-size effect in an initial experiment. In the next experiment, a 100% valid postcue was added to test for an effect of decision. This postcue did not change the set-size effects. In the critical experiments, a 100% valid cue was provided during the retention interval between displays, or only one stimulus was presented in the second display (local recognition). For both of these conditions, there was little or no set-size effect. This pattern of results was found for both hard-to-discriminate stimuli typical of perception experiments and easy-to-discriminate stimuli typical of memory experiments. These results are consistent with capacity limits in memory retrieval, and/or comparison. For these set sizes, the results are not consistent with capacity limits in perception, memory encoding or memory storage.

**Significance Section:** The change detection paradigm is often used to demonstrate effects of divided attention. But it is not clear whether these effects are due to perception, memory, or judgment and decision. In this article, we present new evidence that the divided attention effect in change detection is due to limits in memory retrieval or comparison processes. These results are not consistent with limits in perception, memory encoding or memory storage.

---

One way to study the effects of divided attention is to manipulate set size in the change detection paradigm. Change detection typically consists of presenting two displays of multiple stimuli and asking an observer to judge whether there has been a change between the displays in one or more of the stimuli (e.g. Griffin & Nobre, 2003; Keshvari, van den Berg, & Ma, 2013; Scott-Brown & Orbach, 1998; Woodman, Vogel, & Luck, 2012). Usually, the displays are shown sequentially with a brief separation to prevent transients from signaling the change. For success in this task, the observer must process the sensory input from the first display, encode and store the stimuli in memory, process the second display, retrieve information about the first display, compare the corresponding stimulus representations, and make a decision on the basis of these comparisons. When displays contain multiple stimuli, any decline in performance compared to a single stimulus is called a set-size effect. Such effects have been attributed to a variety of processing stages including perception (Pestilli, Carrasco, Heeger, & Gardner, 2011), memory (e.g. Awh, Barton, & Vogel, 2007; Luck & Vogel, 1997; Rouder, Morey, Cowan, Zwilling, Morey & Pratte, 2008; Wilken & Ma, 2004; Zhang & Luck, 2008) and decision (Scott-Brown, Baker & Orbach, 2000). We pursue the question of which of these processing stages cause set-size effects in change detection.

## Alternative Theories of Change Detection

Consider next a quick review of theories with capacity limits in perception, memory or judgment and decision. In perception, there are many theories that posit limited capacity and predict set-size effects (reviewed in Pashler, 1998; Scharff, Palmer, & Moore, 2011). At one extreme are theories suggesting a bottleneck that allows only one stimulus to be identified at a time (Broadbent, 1958; Lachter, Foster & Ruthruff, 2004). Other theories are less severe suggesting some kind of limited capacity or resource that is divided among relevant stimuli (e.g. Kahneman, 1973; Navon & Gopher, 1979). These theories predict that performance in divided attention tasks is impaired because the stimuli are in competition for the limited capacity in perceptual stages. Finally, there are also theories that assume no capacity limits in the perception of simple features or feature contrast (e.g. Palmer, 1994; Scharff et al., 2011).

Another way that perception can cause set-size effects is from sensory interactions that are largely not attentional. One of the best known such phenomena is crowding (e.g. Bouma, 1970; Pelli, Palomares, & Majaj, 2004). Crowding is a stimulus interaction that limits perception in a variety of tasks (reviewed in Whitney & Levi, 2012). Unless care is taken, increasing set size decreases the spacing between stimuli and increases the degree of crowding. Such crowding effects have also been shown in memory experiments (Tamber-Rosenau, Fintzi, & Marois, 2015). Another phenomena that can be confounded with set size is stimulus heterogeneity. Increasing heterogeneity amplifies set-size effects in both visual search (Rosenholtz, 2001) and visual memory (Lin & Luck, 2009). Finally, changes in stimulus configuration can also affect set-size effects in perception (Nothdurft, 1993) and visual memory (Silvis & Shapiro, 2014).

Theories of memory also assume a variety of capacity limits that can predict set-size effects (see reviews in Brady, Konkle, & Alvarez, 2011; Oberauer, Farrell, Jarrold, & Lewandowsky, 2016). The first display provides a set of study stimuli to be encoded, stored and later retrieved. The second display provides a set of test stimuli that need to be compared to the corresponding study stimuli. Each stage of memory processing has the potential of imposing a capacity limit. For encoding, stimuli might be moved into memory serially (Becker, Miller, & Liu, 2013) or in parallel with capacity limits (Rideaux, Apthorp, & Edwards, 2015; Rideaux & Edwards, 2016). Storage limits might exist in terms of the number of objects (Luck & Vogel, 1997) or of the quality of the representations (e.g. Keshvari et al., 2013). For retrieval, some theories propose serial (McElree & Dosher, 1993) or parallel (McElree & Dosher, 1989) access, or an effect of interference during retrieval that depends on the number of relevant stimuli (Oberauer & Lin, 2016). A yet longer list of possibilities is described in the General Discussion. Each of these broad hypotheses - limited capacity in encoding, storage or retrieval - can predict a set-size effect in the change detection task.

Following memory, the final stages of processing are judgement and decision and they too can cause set-size effects. At this point in processing, observers already have the relevant information in memory and need to compare this information in order to make a response. We divide the judgment and decision processes into two parts. First, there might be a capacity limit in the comparison process between the two displays (Angelone, Levin, & Simons, 2003; Farell, 1985; Fernandez-Duque & Thornton, 2000; Hyun, Woodman, Vogel, Hollingworth, & Luck, 2009; Mitroff, Simons, & Levin, 2004; Simons, Chabris, Schnur, & Levin, 2002). Second, there might be a capacity limit due to noise compounded from having to combine multiple stimulus comparisons in the final decision (Palmer, Verghese, & Pavel, 2000; Sperling & Dosher, 1986; Scott-Brown, et al., 2000; Tanner, 1961).

## Our Study

Our overarching goal is to find the locus of set-size effects. We also have three supporting subgoals that guide our investigation. In typical set-size experiments, set sizes are varied over a range of 1, 2, 4, and more stimuli. In contrast, our first subgoal is measure the effect of increasing set size from just 1 to 2 stimuli because these effects reveal the *initial source* of capacity limits (e.g. Bae & Flombaum, 2013; White, Palmer, & Boynton, 2018; Williams, Hong, Kang, Carlisle, & Woodman, 2013). Effects of higher set sizes are expected to share these initial capacity limits as well as possibly adding capacity limits imposed by other processes.

Our second subgoal is to measure set-size effects that are purely attentional and not due to nonattentional phenomena such as crowding (Tamber-Rosenau et al., 2015), stimulus heterogeneity (Lin & Luck, 2009), and other configural phenomena between stimuli (Silvis & Shapiro, 2014). To do this, rather than manipulate set size, we manipulate the number of relevant stimuli using a 100% valid precue (e.g. Makovski, Sussman, & Jiang, 2008; Wright & Green, 2000). Manipulating relevant set size allows the visual displays to be identical in all conditions and thus holds constant any stimulus-driven effect (Palmer, 1994). This *effect of relevant set size* give the best chance of measuring purely attentional effects.

Our third subgoal is to conduct an experiment relevant to research in both perception and memory. Change detection in memory research primarily differs from its use in perception research by the choice of stimuli. Rather than using hard-to-discriminate stimuli such as low-contrast Gabors, or small luminance changes, memory research typically uses easy-to-discriminate stimuli such as high-contrast bars, colored patches or nameable objects (e.g. Hollingworth, 2003; Luck & Vogel, 1997; Wilken & Ma, 2004). To address both bodies of research, we used both hard-to-discriminate stimuli and easy-to-discriminable stimuli. To give away one aspect of our results, similar effects were found for both hard- and easy-to-discriminate stimuli. Thus, for our measurements, these two experimental traditions gave a common result.

The locus of the set-size effects can be revealed by cues during different points in the processing sequence. Figure 1 shows a schematic of the processing stages that are associated with hypotheses for capacity limits in change detection. The arrows show information about two stimuli moving through each stage. The hypotheses for different loci of capacity limits can be distinguished by the use of cueing. Consider first a *retention-interval cue* (often called a retro-cue) between the two arrays (reviewed in Souza & Oberauer, 2016). It can affect memory maintenance, memory transfer, memory retrieval, comparison and decision processes but not perception, memory encoding and the initial memory storage. This is because perception, memory encoding and initial storage are already complete when the cue is presented (Beck & van Lamsweerde, 2011; Griffin & Nobre, 2003; Hollingworth & Maxcey-Richard, 2013; Landman, Spekreijse, & Lamme, 2003; Makovski et al., 2008). In the schematic, the retention-interval cue appears between the encoding-and-storage stage, and the retrieval stage. After the cue, only one stimulus representation need be retrieved and compared. Alternatively, consider a *postcue* that follows the second stimulus display. It does not influence memory or comparison, but can affect the final decision processes (e.g. Beck & van Lamsweerde, 2011; Hawkins et al., 1990). Thus, a retention-interval cue and a postcue eliminate the need for processing multiple stimuli at different points in the processing sequence.

**Figure 1.**
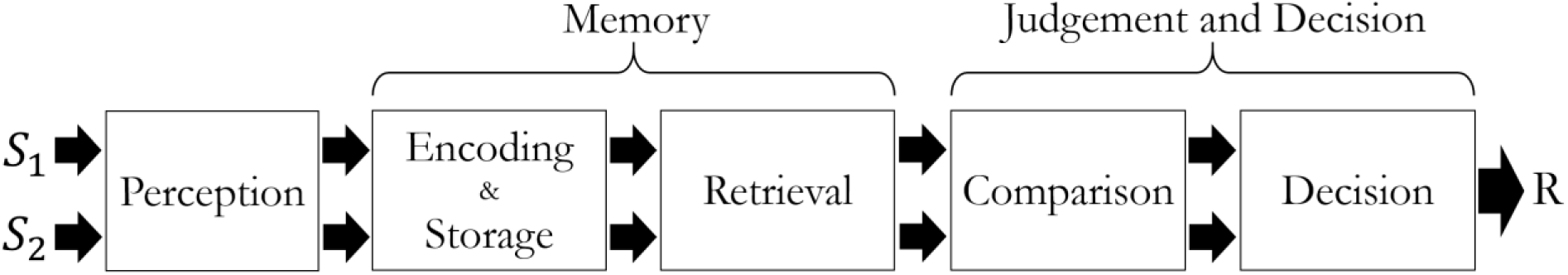
Schematic of the stages of processing necessary for the change detection task. Each stage is associated with one or more potential capacity limits.

## Overview of Experiments

In our first experiment, a basic form of change detection was used in which two oriented Gabor stimuli were presented in the first display, followed by a blank, and then a second pair of Gabor stimuli. The observer’s task was to report whether either Gabor changed in orientation from the first to the second display. The decision in this task has a many-to-one mapping because a change at either location maps to the same response. This means that any noise in the stimulus representations is compounded across sides and may obfuscate effects occurring in earlier processing stages. To remove this contribution, in the second experiment, a 100% valid postcue and independence across sides is introduced to make the stimulus response mapping one-to-one. This tests for a contribution from the final decision process to the set-size effects.

In the third experiment, we modify the procedure with the addition of a 100% valid retention-interval cue to test the contribution of capacity limits on memory maintenance, retrieval and comparison. If a process such as memory encoding or the available storage limits performance, the set-size effect with the modified procedure should remain the same as in the first two experiments because nothing about the displays or stimuli have changed up to this point. If, however, change detection is limited by memory maintenance, retrieval or comparison, the set-size effect should be reduced, or in the extreme, eliminated. In the fourth experiment, the results are further extended using a local recognition task to distinguish memory maintenance from retrieval and comparison. In summary, these experiments allow one to narrow the possible loci of capacity limits in change detection.

We conducted all of the experiments in two ways, first low-contrast Gabor patches were used as hard-to-discriminate stimuli as common in perceptual experiments. The contrast was chosen so that the mean performance with a single relevant stimulus was about 80% correct. In addition, high-contrast Gabor patches were used as easy-to-discriminable stimuli as common in memory experiments. Under these high-contrast conditions, mean performance with a single relevant stimulus was about 98% correct.

## General Methods

### Observers

Twelve observers participated in each of the experiments. All observers had normal or correct-to-normal vision. One of the observers was author JM. All observers (except JM) were compensated $20/hour. All observers gave written and informed consent in accord with the human observers Institutional Review Board at the University of Washington, in adherence with the Declaration of Helsinki.

To determine the minimum number of observers needed to detect a set-size effect, we conducted a power analysis based on pilot data from a previous Gabor detection experiment. In the previous experiment, observers (*N* = 5) each completed 1920 trials in a simple Gabor detection experiment comparing detection at one versus two possible cued locations (relevant set sizes 1 versus 2). The stimuli and procedure were similar to the present experiments. The observed set-size effect was 4.2% ± 1.1%. The standard deviation of this effect across observers was 2.4%. A power analysis was done for a yet smaller set-size effect of 2%. Using a paired sample, one-tailed t-test and a power of 80%, the minimum number of observers required was 11. For good measure, we chose to use 12 observers in each experiment.

### Stimuli and Procedure

In all four experiments, the basic task was to detect whether the orientation of a Gabor in noise changed from a first display to a second display. Figure 2 shows a schematic of the procedure for each of the conditions of Experiment 1. Trials were blocked by condition: relevant set size 1 (left and right), and relevant set size 2. For all conditions, observers began by foveating a fixation cross at the center of a gray screen (500 ms; 50% of max luminance). This was followed by a 100% valid precue consisting of two lines on either side of the fixation cross (1° eccentricity; 500 ms). For relevant set size 1, the lines were different colors (red and blue); for relevant set size 2, the lines were the same color. Each observer was allocated a cue color that indicated the relevant side (colors were counterbalanced across observers). An earlier version of the experiments did not have a precue with set size 2. However, there was no difference in the results for observers who ran under these conditions so data was collapsed for analysis. Following the precue, a display containing two patches (6° x 6°) of dynamic noise each with a Gabor that appeared on either side of fixation, centered at 4° eccentricity on the horizontal meridian. After the first display (1000 ms) there was a delay with only the fixation cross (1000 ms) followed by a second display that contains two noise patches each with a Gabor (1000 ms). After a brief delay with the fixation cross alone (250 ms) a 100% valid postcue appeared until the observer responded whether or not the orientation of either cued Gabor changed from the first to the second display interval. For Experiment 1, the postcue was identical to the precue. Only one response was required. Responses were given on a rating scale (likely-no, guess-no, guess-yes, likely-yes) to measure an ROC curve. Auditory feedback was provided for incorrect responses (180Hz).

**Figure 2.**
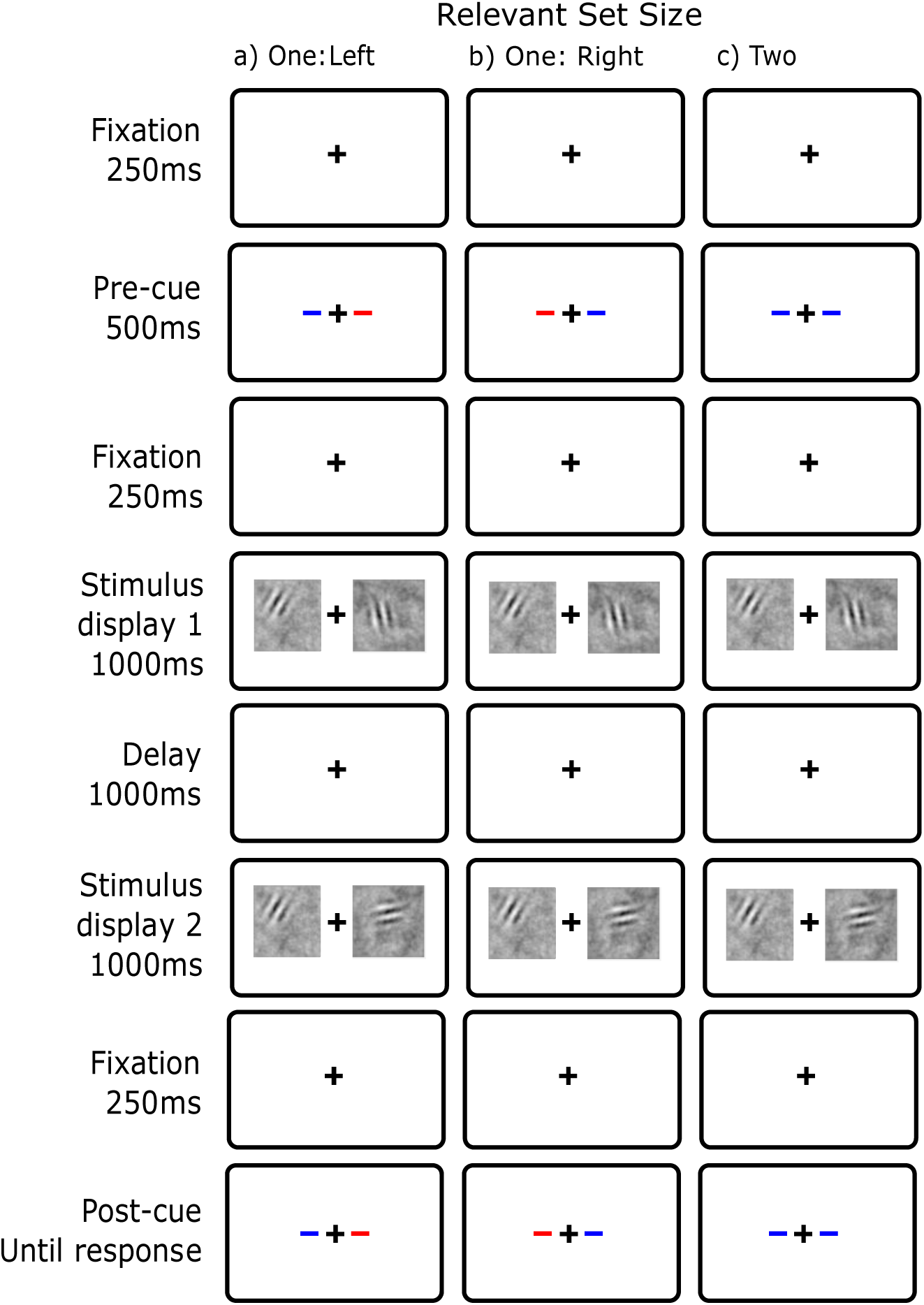
Display conditions for basic change detection (Experiment 1) with blue as the cue color. a) Relevant Set Size 1 - Left b) Relevant Set Size 1 - Right c) Relevant Set Size 2. Unlike this illustration, all experiments had the entire screen a middle gray.

Each block consisted of 24 trials from one of the three conditions: set size 2, set size 1 left, and set size 1 right. A single experimental session included four set-size-2 blocks, two set-size-1-left blocks and two set-size-1-right blocks. Each observer performed practice sessions in which the Gabor contrast was lowered gradually until performance was stable around 80% for set size 1. Observers then completed 10 sessions at this near threshold contrast resulting in 1920 trials overall per observer. Each session took 20-25 minutes, and typically two sessions were run back-to-back within an hour. We also collected four sessions with high-contrast Gabors (80% contrast) from each observer to assess performance with highly visible stimuli. Inadvertently, two observers in Experiment 1 did not complete 1 or 2 low-contrast sessions, and two observers in Experiment 2 did not complete 2 high-contrast sessions.

#### Noise Movies

The “movies” had *1/f* noise in space and time and played for 1000 ms with an effective framerate of 30 Hz. The movies were generated as follows: Each frame was first populated with independent Gaussian noise at each pixel, with zero mean and unit variance. The frame was then filtered using a 2D Fourier transform such that the amplitude of each spatial frequency component *fs* was proportional to *1/fs*. Then, the whole movie was similarly filtered in time so that the amplitude of each temporal frequency *ft* was proportional to *1/ft*. The pixel values were then rescaled to have a standard deviation of 0.12 (a relatively low luminance contrast). The local contrast of each frame was attenuated at the edges by a linear ramp down to zero beginning 0.5° from the nearest edge.

#### Gabors

The Gabor patches had spatial frequency of 1 cycle/° and were windowed by a 2D Gaussian with a standard deviation of 0.5° and truncated to a total width of 2°. The Gabor could appear anywhere within the noise image, as long as the edges of the truncated width were at least 0.5° from the edges of the noise. The Gabor’s contrast was modulated in time by a Gaussian envelope (standard deviation 50 ms). The time of maximal contrast was chosen from a uniform distribution, excluding the first and last 200 ms of the movie, but constrained to appear at the same time on both sides of the stimulus display to avoid the possible advantage of an attention switching strategy. Orientations were drawn uniformly from two sets of non-overlapping standards [11.25°, 56.25°, 101.25°, 146.25°] and [33.75°, 78.75°, 123.75°, 168.75°]. The standards were offset so that the same orientation was never present on both sides at once. The set of values used for each side randomly varied so no orientation was associated with a side.

On all trials, eye position was recorded using an Eyelink II, 2.11 with 250 hz sampling (SR research, ON). The position of the right eye was recorded for all trials, and trials were included for analysis only if fixation was confirmed. When fixation failed, observers were alerted with five consecutive high frequency tones and the trial aborted. The percentage of aborted trials for each observer in each experiment ranged from 1.7% to 14% with an overall mean including all experiments of 5.7 ± 0.8%.

### Analysis

Observers responded with one of four key presses that indicated likely-no, guess-no, guess-yes, or likely-yes. These ratings were used to form a receiver operating characteristic (ROC) function and performance was summarized as the percent area, *A’*, under the ROC function. *A’* is equivalent to the percent correct measured by forced-choice paradigms (Green & Swets, 1966). To estimate *A’* the trapezoid method was used to avoid making distributional assumptions (Macmillan & Creelman, 2004) and converted to a percentage. The difference in *A’* between set size 2 and 1 is our primary measure of the effect of divided attention. We refer to it as the *relevant-set-size effect*.

### Experiment 1: Basic Change Detection

Our first experiment was designed to estimate the magnitude of the set-size effect in a version of change detection that is typical of the literature (e.g. Keshvari, van den Berg, & Ma, 2013). This task consists of two stimulus displays separated by a blank. In each display, there was a stimulus on both sides of fixation. For set size 1, if a change occurs it was restricted to the precued side whereas for set size 2, the change can occur on either side and the observer must make a single decision for the whole display. Given that the task is made up of two possible events that can map to the same response (a many-to-one mapping) this is sometimes called a compound task (Sperling & Dosher, 1986) and is commonly used in visual search. No set-size effect is expected if all processing stages have unlimited capacity (perception, memory and decision).

#### Methods

The method was as described in the General Methods section. The specific task is shown in Figure 2 and proceeded as follows. The first and second stimulus displays contained a Gabor in both the left and the right side. On 50% of trials a change in orientation of 90° occurred on one of the relevant sides. In the relevant-set-size-1 blocks, the change could occur on only the precued side, and the uncued side always remained unchanged in orientation. In relevant-set-size-2 blocks, the change could occur on either side but not both. The observer’s task was to make a yes-no response as to whether a change had occurred anywhere.

#### Results

The effects of relevant set size on accuracy (collapsed across sides) are shown in Figure 3a. Consider first the low-contrast conditions shown by the open symbols. Performance was better for relevant set size 1 (Mean = 80.1, SE = 0.9) than set size 2 (Mean = 72.2, SE = 1.1). This is a difference of 7.9%, 95% CI [5.7, 10.1], t(11) = 7.91, *p* < .001.

**Figure 3.**
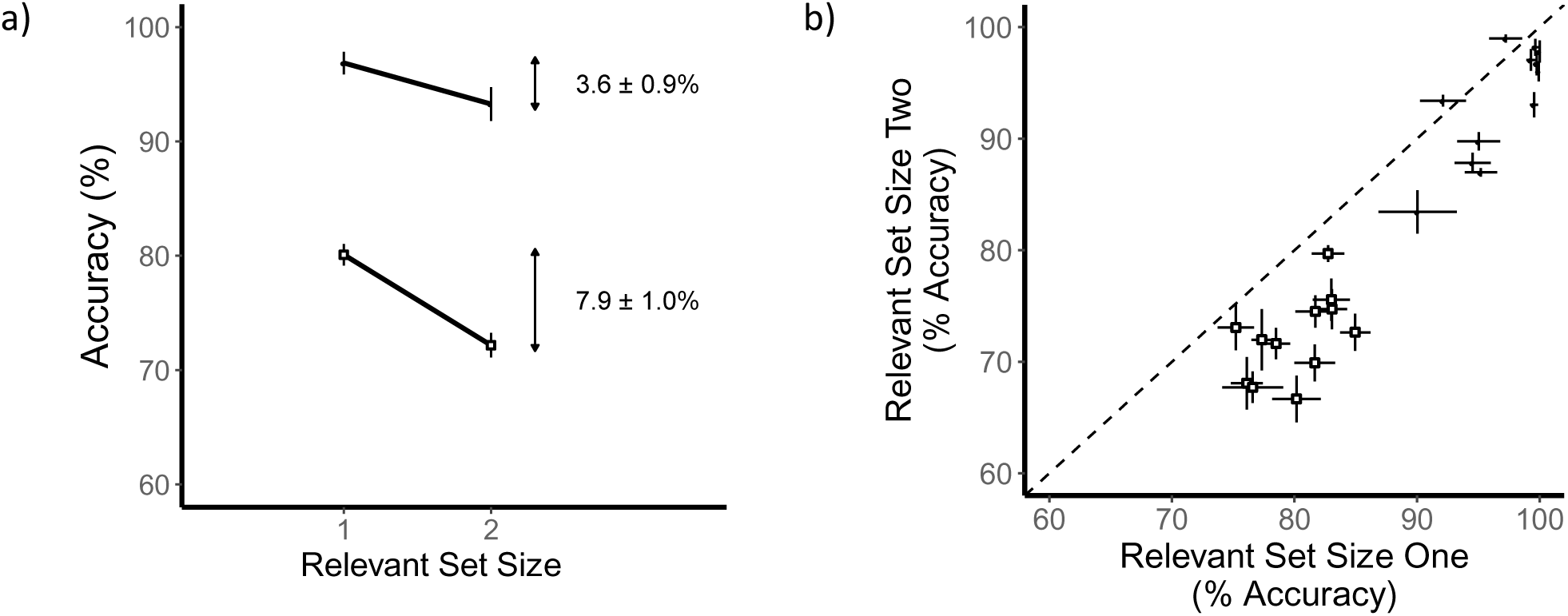
Results for basic change detection (Experiment 1). Low-contrast conditions are shown by open squares and high-contrast conditions are shown by filled circles. a) A graph of performance for relevant set sizes 1 and 2. There are reliable set-size effects for both contrast conditions. b) A scatterplot of individual observer performance for set size 2 versus set size 1. All but two observers show the expected set-size effect. Error bars are standard error of the mean.

Now consider the high-contrast conditions that are shown by the filled symbols. There was also a significant effect of relevant set size. Performance was better for set size 1 (Mean = 96.9, SE = 1.0) than for set size 2 (Mean = 93.3, SE = 1.5). This is a difference of 3.6%, 95% CI [1.5, 5.6], t(11) = 3.88, *p* = .003.

These effects of relevant set size were found for almost all observers. In Figure 3b, a scatterplot of individual observers is shown with the accuracy in set size 2 plotted against the accuracy in set size 1. Points falling below the identity line indicate worse performance with the larger set size.

#### Discussion

The results of the basic change detection experiment are consistent with similar studies in showing a set-size effect (Keshvari et al., 2013; Luck & Vogel, 1997; Scott-Brown & Orbach, 1998). Moreover, it is consistent with prior studies showing such an effect for set sizes 1 versus 2 (Bae & Flombaum, 2013; Williams, et al., 2013). Thus, even for two stimuli, one or more component processes must be limiting performance with multiple stimuli.

### Experiment 2: Postcues

In the next experiment, we addressed the role of decision in change detection. Simple change detection as in Experiment 1 includes dependencies across sides because different events can lead to the same response (e.g. a change on the left, or right will lead to a change response). This many-to-one mapping complicates the interpretation of the results because it obfuscates the source of information used in the decision (Braun & Julesz, 1998; Shaw, 1980; Sperling & Dosher, 1986). In the next experiment, each stimulus judgement was made an independent task (sometimes called a concurrent task or dual-task; Sperling & Dosher, 1986), and a postcue was used to indicate the relevant stimulus and response. The task now has a one-to-one mapping between stimulus and response (illustrated in the schematic in Figure 4). If the result of Experiment 1 is due to only the effect of the compounded decision error then the effect should be eliminated in Experiment 2. Furthermore, the stimulus displays for relevant set size 1 can now be identical to relevant set size 2. Changes occur on both sides independently with 50% probability. In this experiment, the precue is the only difference between the conditions.

**Figure 4.**
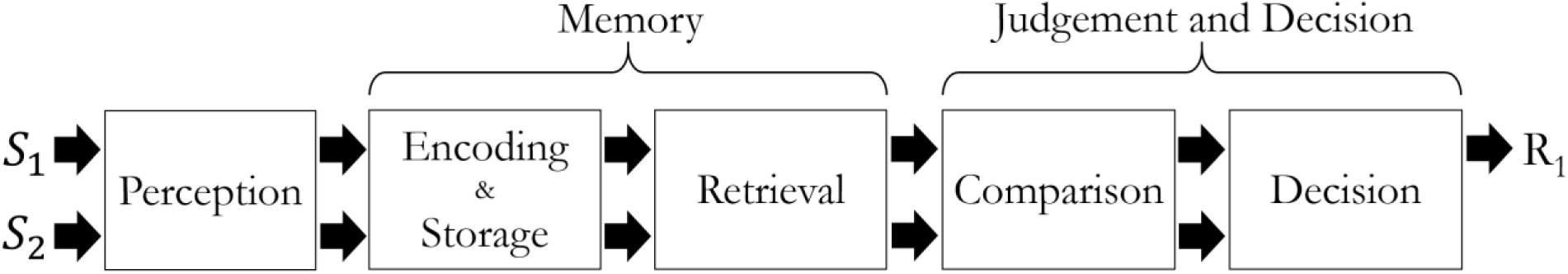
Schematic of the stages of processing for the change detection task with a late postcue (Experiment 2). For the example trial shown, a response is required for only the first stimulus (S1).

#### Methods

The General Methods were used except that (a) the presence of a change is independent on the left and right side - changes occur on one or both sides in both of the set-size conditions, and (b) observers must respond to whether a change occurred within the postcued side. The observer used two independent sets of response keys corresponding to the left and right side (but only one response is made on each trial).

**Figure 5.**
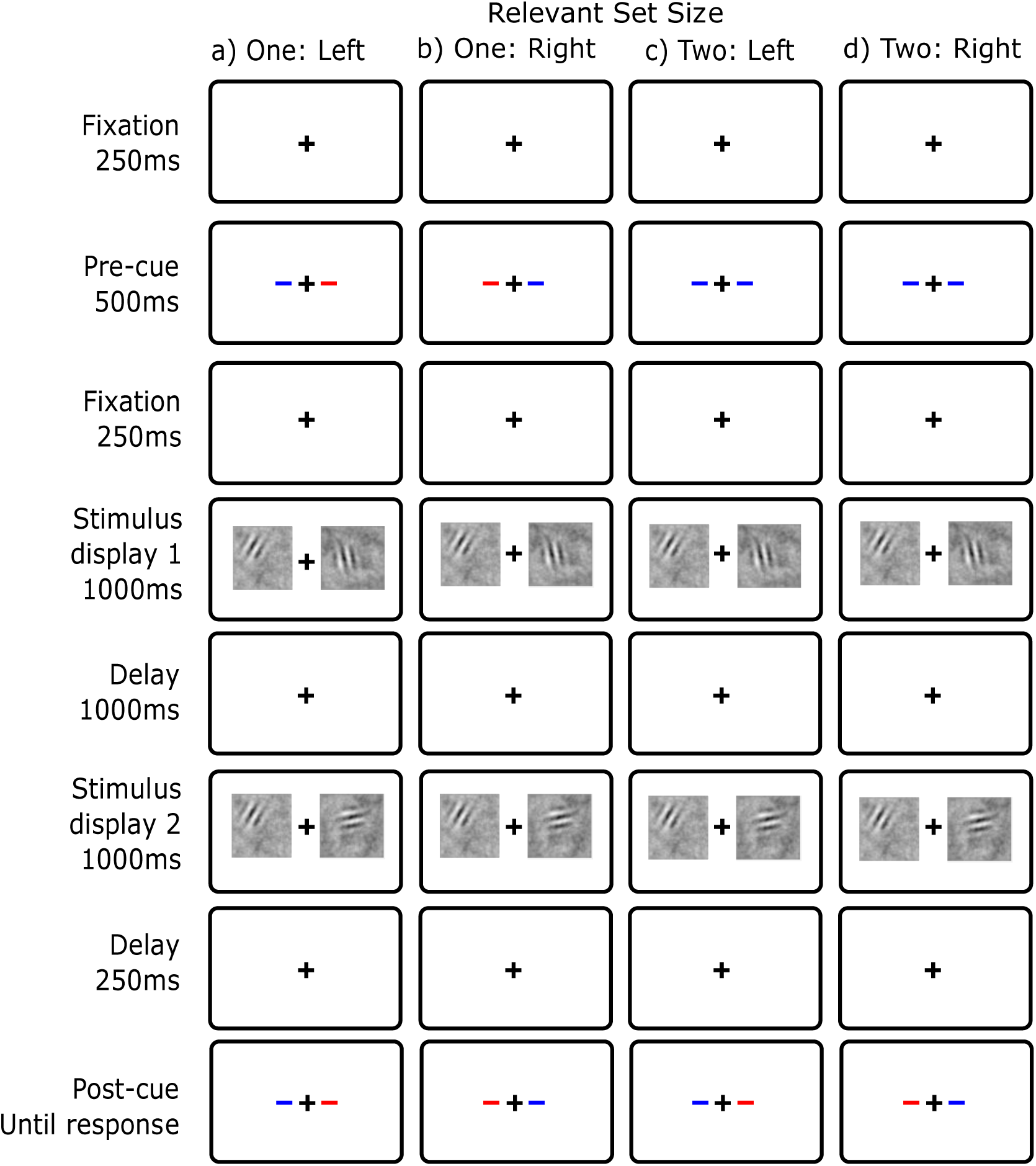
Display conditions for change detection with a late postcue (Experiment 2). Blue is the cue color. a) Relevant Set Size 1 - Left: The left side is precued with 100% validity as the side that will be tested on this trial. b) Relevant Set Size 1 - Right: The right side is precued as relevant. c) Relevant Set Size 2 - Left: Both sides were precued as potential response sides and the left side is later postcued for response. d) Relevant Set Size 2 - Right: Both sides were precued as relevant and the right side is later postcued for response.

#### Results

The effect of relevant set size on accuracy is shown in Figure 6a. For the low-contrast condition, performance is better for set size 1 (Mean = 82.1%, SE = 1.3%) than for set size 2 (Mean = 75.0%, SE = 1.7%). This is a difference of 7.1%, 95% CI [4.1, 10.1], *t*(11) = 5.22, *p* < .001. For the high-contrast condition, performance is also better for set size 1 (Mean = 98.1, SE = 0.6) than set size 2 (Mean = 91.0, SE = 1.8). This is a difference of 7.2%, 95% CI [3.9, 10.4], t(11) = 4.9, *p* < .001. The results for each individual observer are shown in Figure 6b. The set-size effect is found for almost every observer.

**Figure 6.**
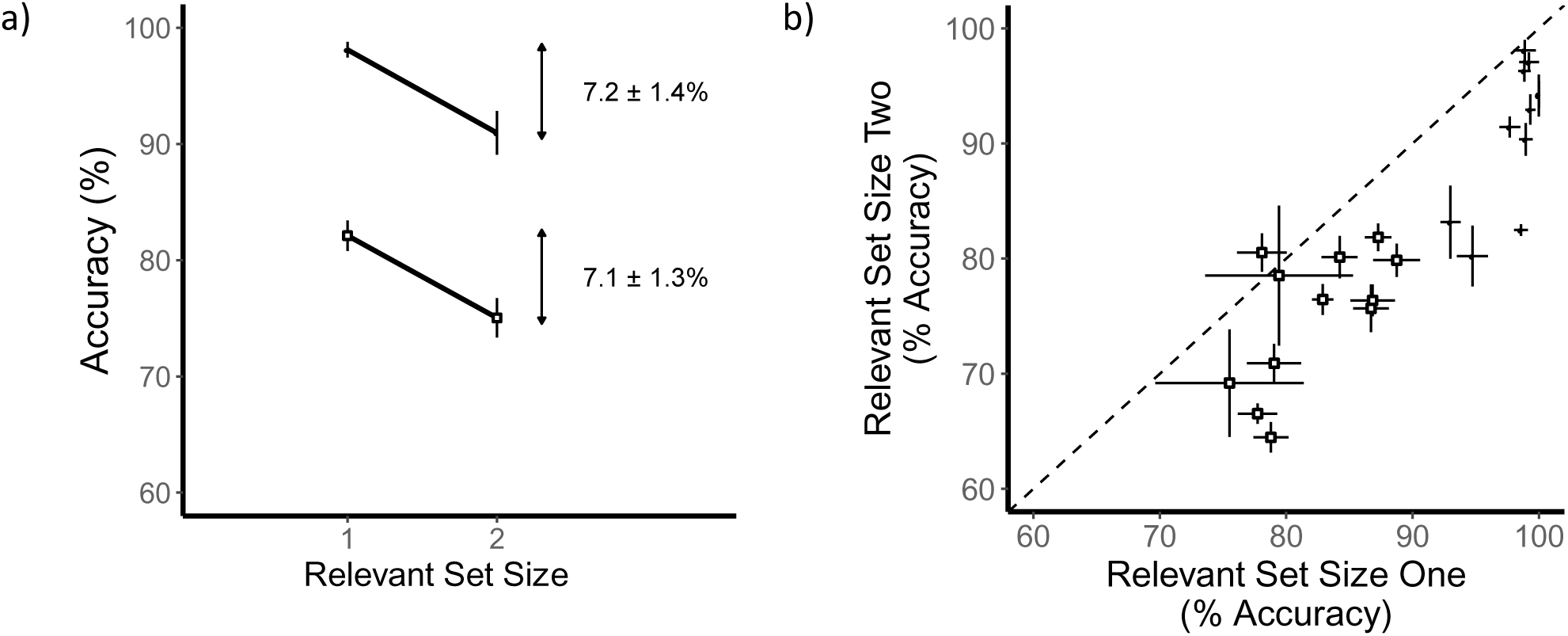
Results for change detection with a postcue (Experiment 2). Low-contrast conditions are shown by open squares and high-contrast conditions are shown by filled circles. a) Accuracy is shown for relevant set sizes 1 and 2. There are reliable effects for both contrast conditions. b) A scatterplot of individual observer performance for set size 2 versus set size 1. All but one observer shows the expected set-size effect.

#### Congruency

The results of this experiment can be further analyzed by the congruency of the stimulus events at each location on each trial. Congruent trials have the same stimuli event (e.g. a change) occurring at both locations. Incongruent trials have different stimuli events occurring at each location (e.g. a change on one side, and not on the other). Effects of congruency are evidence of interactive processing of the two stimuli (e.g. Navon & Miller, 1987). However, in this and the following experiments, there were no congruency effects on performance (see Appendix). Thus, there is no sign of interactive processing.

#### Orientation similarity and perceptual grouping

When two orientations are presented together, sometimes they can form a single perceptual representation or *group* (Silvis & Shapiro, 2014). If such perceptual grouping occurs for our observers, it undermines our assumption of testing one versus two stimuli. Due to our stimulus design, there were never identical orientations on both sides in a given stimulus display. However, there are still pairings that might be grouped into either corners or almost parallel lines. Despite this possibility, there was no evidence of perceptual grouping (see Appendix).

#### Discussion

In this study, the postcue did not reduce set-size effects. This is similar to some previous studies (e.g. Luck & Vogel, 1997; Wheeler & Treisman, 2002) but others have found that set-size effects are reduced by the postcue (e.g. Beck & van Lamsweerde, 2011; Hollingworth, 2003). This literature is examined more closely in the discussion. In sum, decision appears to not contribute to the set-size effect in this task.

### Experiment 3: Retention-interval cue

In Experiment 3, we begin to consider whether effects before memory retrieval contribute to the set-size effect. This experiment is identical to Experiment 2 with the postcue, except that a retention-interval cue (e.g. Griffin & Nobre, 2003) was inserted between the two stimulus displays. The retention-interval cue indicated the relevant stimulus (i.e. it matched the postcue). This task therefore required observers to retrieve from memory only the orientation on the relevant side, and make a single comparison decision on the relevant side before responding. This is illustrated in the schematic in Figure 7. Thus, it should eliminate the effect of capacity limits in retrieval or comparison.

**Figure 7.**
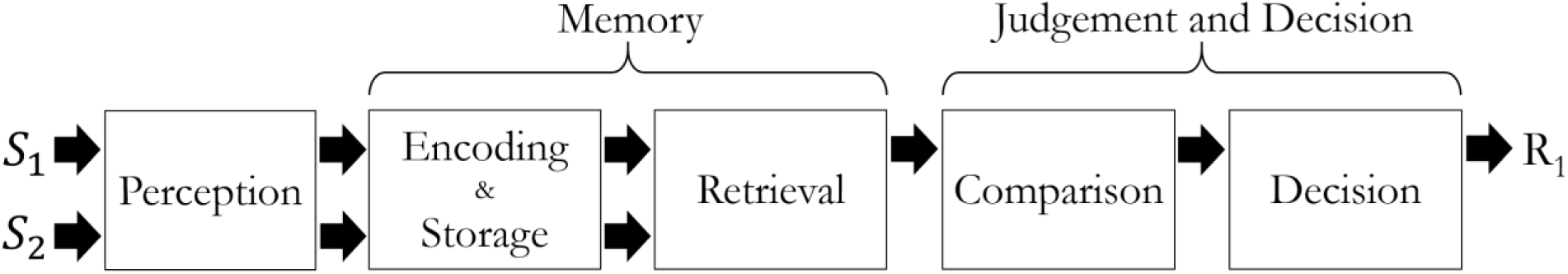
Schematic of the stages of processing for the change detection with both a postcue and a retention-interval cue (Experiment 3). For the example trial shown, a retention-interval cue indicates that only the first stimulus (S1) is relevant. Thus, there is no continued processing for the uncued side.

Observers still had to perceive and encode the two stimuli from the first stimulus display, and store them in memory until the retention-interval cue. Thus, Experiment 3 has the same perception, memory encoding and storage requirements as Experiment 2, but different memory retrieval and comparison demands. If perception, memory encoding, or memory storage of two orientations is the limiting factor then the set-size effect should persist with a retention-interval cue. If, however, memory retrieval or comparison is the limiting factor then the set-size effect will be eliminated.

There are two versions of the storage hypothesis that make different predictions than a limit based simply on storage capacity. The first we call *selective maintenance*. By this hypothesis, there are maintenance processes such as rehearsal during the retention interval. The retention-interval cue allows for selective maintenance of the relevant stimulus for the part of the retention interval following the cue. Thus, this hypothesis predicts the retention-interval cue can reduce the set-size effect.

The second hypothesis we call *selective transfer*. This idea depends on the existence of multiple memory stores. For example, consider the possible role of both an iconic memory store and a short term store. A retention-interval cue that was presented while the iconic memory store was still available would allow the selective transfer of information about the relevant stimulus. For our experiment the timing in the presence of dynamic noise make the persistence of iconic memory unlikely (but see the Appendix for further evidence). A more plausible multiple store model was presented by Sligte, Scholte, & Lamme (2008). They propose that the relevant visual memory consists of both a high capacity and fragile store (*Fragile VSTM*) and a lower capacity and durable store (*Traditional VSTM*). They propose that fragile VSTM is durable enough to last a few seconds under some conditions. As a result the retention-interval cue could allow the selective transfer of information about the relevant stimulus into the more durable memory store. This predicts the retention-interval cue will reduce the set-size effect.

In summary, different storage hypotheses make different predictions about the effect of the retention-interval cue. The storage capacity hypothesis predicts no effect. In contrast, the selective maintenance and selective transfer hypotheses do predict an effect of the retention-interval cue.

#### Methods

The stimuli in this experiment were identical to Experiment 2 except for the additional retention-interval cue. The retention-interval cue was always identical to the postcue. For relevant set size 1, this new cue provided no additional information. For relevant set size 2, the observer knew which side was relevant during the retention interval.

**Figure 8.**
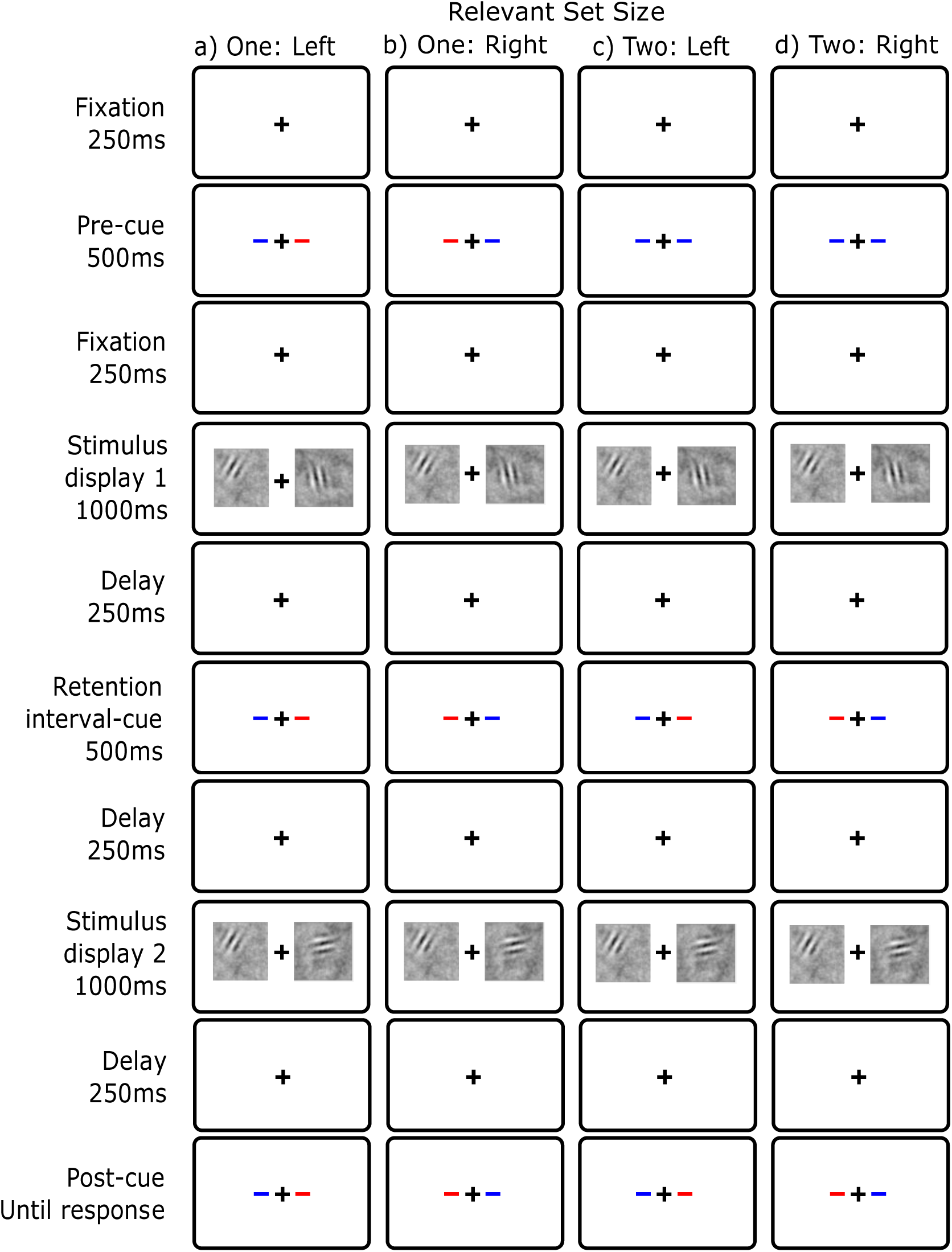
Display conditions of change detection with a retention-interval cue (Experiment 3). Blue is the cue color. The retention-interval cue appears between the two displays and matches the postcue. a) Relevant Set Size 1 - Left, b) Relevant Set Size 1 - Right, c) Relevant Set Size 1 - Left, d) Relevant Set Size 1 - Right.

#### Results

The effect of set size is shown in Figure 9a. For the low-contrast condition, there was no reliable difference between set size 1 (M = 79.3, SE = 1.1) and set size 2 (M = 78.5, SE = 0.9) conditions. This is a difference of 0.8%, 95% CI [-1.0, 2.5], t(11) = 0.97, *p* = .35. For the high-contrast condition, there was also no reliable difference (set size 1, M = 98.0, SE = 0.6; set size 2, M = 97.0, SE = 1.0, difference = 1.1, 95% CI [-0.2, 2.3], t(11) = 1.9, *p* = .10. The lack of set-size effects is also clear in the scatterplot of individual observers (Figure 9b).

**Figure 9.**
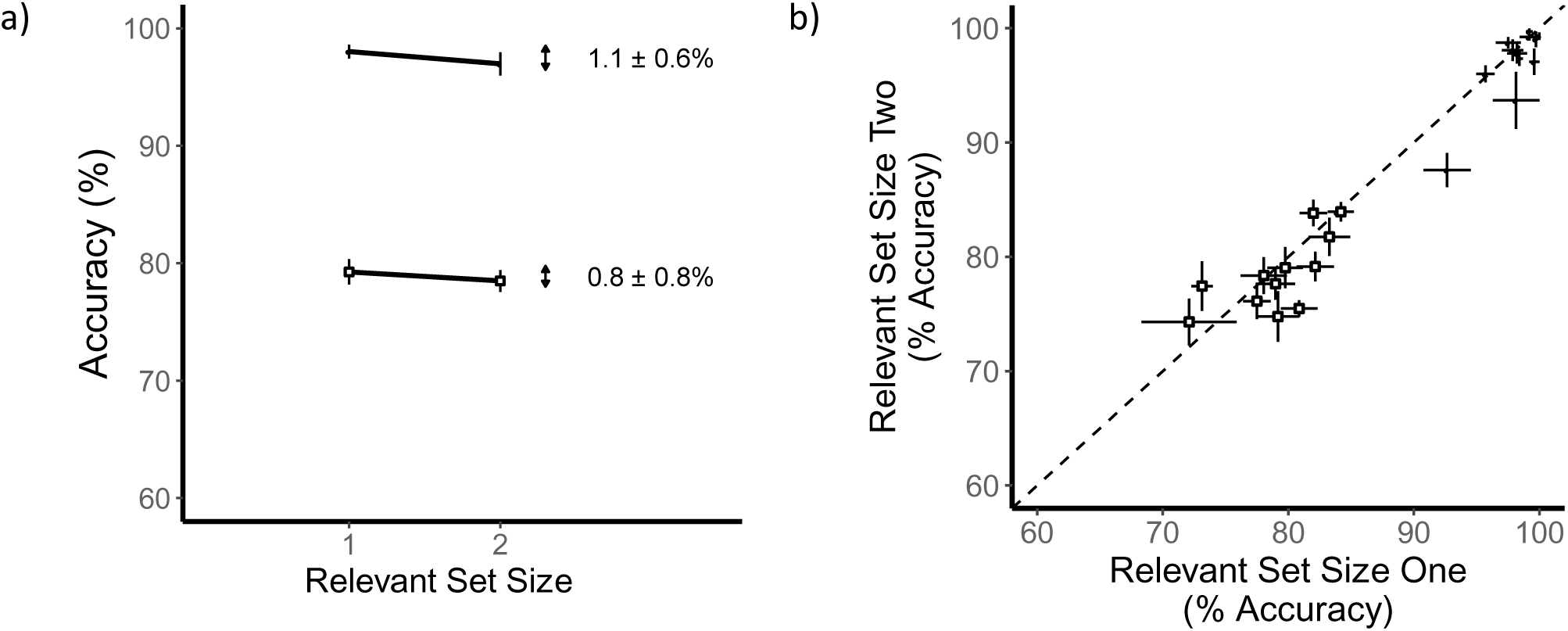
Results for change detection with a retention-interval cue (Experiment 3). Low-contrast conditions are shown by open squares and high-contrast conditions are shown by filled circles. a) Graph of accuracy as a function of relevant set size. b) Scatterplot of individual observer performance for set size 2 versus set size 1. There is no reliable effect of set size.

#### Discussion

The lack of a set-size effect in Experiment 3 suggests that there is no capacity limit up to and including the initial storage of the stimuli. In other words, for two simple stimuli, there is no limit in perception or for memory encoding and initial storage. Instead, the limiting process must be one of the later storage processes (selective maintenance or selective transfer), retrieval or comparison.

### Experiment 4: Local Recognition

In Experiment 4, we use a local recognition task: The first stimuli are presented on both sides followed by a test display with only one stimulus presented on the response side (Figure 10). This is similar to Experiment 3 in that the observer must make only one memory retrieval and comparison. The new feature of this experiment is that now hypotheses such as selective maintenance and selective transfer no longer predict a reduction in the set-size effect. The information indicating the relevant stimulus comes after the retention interval as part of the test display. If this local recognition task eliminates the set-size effect, it would be consistent with capacity limits in either retrieval or comparison. This paradigm is similar to experiments where a cue is presented simultaneously with the second (or test) stimulus display (Luck & Vogel, 1997; Makovski, Sussman, & Jiang, 2008; Wheeler & Treisman, 2002).

**Figure 10.**
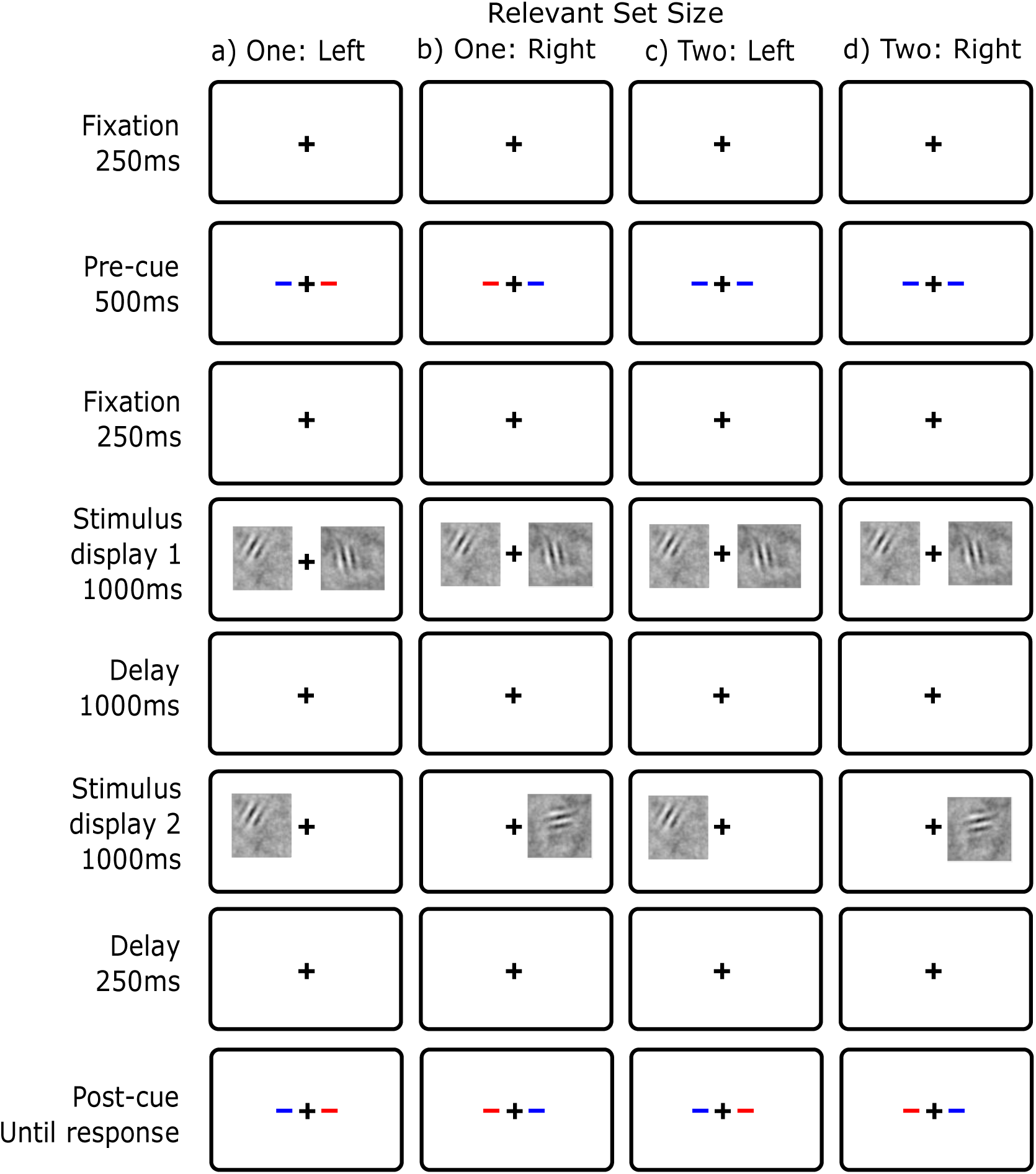
Display conditions for change detection with local recognition (Experiment 4). Blue as the cue color. a) Relevant Set Size 1 - Left, b) Relevant Set Size 1 - Right, c) Relevant Set Size 1 - Left, d) Relevant Set Size 1 - Right.

#### Methods

The stimuli were identical to Experiment 2 except the second stimulus display contained a stimulus on only the relevant side (Figure 10).

#### Results

The effect of relevant set size is shown in Figure 11a. For the low-contrast condition, there was no reliable difference between set size 1 (M = 82.5, SE = 1.2) and set size 2 (M = 80.8, SE = 1.5) conditions. This is a difference of 1.7% 95% CI [-0.1, 3.46], t(11) = 2.08, *p* = .06.

For the high-contrast condition, there was a small but significant effect (set size 1, M = 97.7, SE = 0.8; set size 2, M = 96.5, SE = 1.1, difference = 1.2, 95% CI [0.4, 2.0], t(11) = 3.4, *p* = .006). This small effect is at the limit of what can be measured in these experiments and is a fraction of the effects measured in the basic and postcue experiments. Furthermore, compared to Experiment 3, this effect is reliable because the sample standard deviation is relatively small rather than the difference in accuracy being large. Thus, we emphasize the small size of this effect in contrast with the previous experiments rather than its reliable difference from zero.

**Figure 11.**
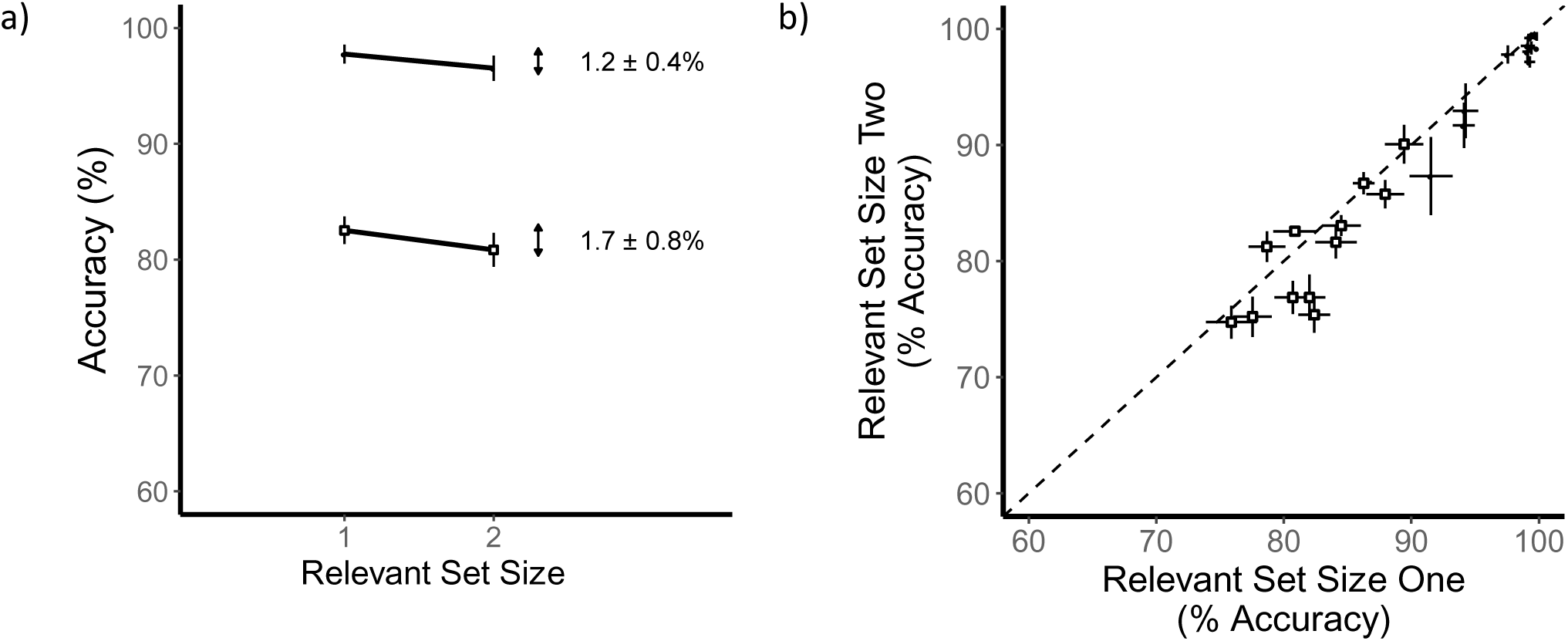
Results for change detection with local recognition (Experiment 4). Low-contrast conditions are shown by open squares and high-contrast conditions are shown by filled circles. a) A graph of accuracy as a function of relevant set size. b) Scatterplot of individual observer performance for set size 2 versus set size 1. The effects of set size are quite small which is similar to Experiment 3.

#### Discussion

In Experiment 4, local recognition showed a smaller set-size effect than change detection in which the second display included all of the stimuli rather than just one. The previous literature on local recognition is mixed. Wheeler & Treisman (2002) investigated change detection in experiments with whole displays and with the single displays of local recognition. They found better performance with local recognition but whether set-size effects were reduced is less clear. In contrast, Jiang, Olson, & Chun (2000) compare these conditions with larger set sizes and found worse performance with local recognition. They attributed this effect to the loss of configural information with the single display in local recognition. The current study minimize the role of configural information, so that might explain why our results were more like those found by Wheeler and Treisman.

The results of Experiment 4 are consistent with the results of Experiment 3 showing that when only one retrieval and comparison must be made there is little or no set-size effect. An alternative explanation for Experiment 3 was that the retention-interval cue between the stimulus displays changed the storage processing in some way. For example, the cue might have allowed selective maintenance of the relevant stimulus information or allowed selective transfer of the relevant stimulus information to a more durable memory. Finding the same result for Experiment 4, which did not have the retention-interval cue, rules out an explanation based solely on a difference in storage processes (Williams & Woodman, 2012; Zhang & Luck, 2008). Instead, the only hypotheses consistent with all experiments involve retrieval and/or comparison.

## General Discussion

In this study of change detection, our goal was to find the locus of capacity limits with just two stimuli. Comparing one versus two relevant stimuli reveals the initial and most restrictive limits on processing. In addition, we manipulated relevant set size to measure purely attentional effects, and measured both easy-to-discriminate and hard-to-discriminate stimuli to address typical experiments for both perception and memory.

### Summary of Results

We measured effects of relevant set size in four kinds of change detection and the results are summarized in Figure 12. Consider first the results for low-contrast stimuli typical of perceptual experiments that are shown in Panel A. For basic and postcued change detection, there was a large set-size effect: performance was better for one relevant stimulus compared to two. These two tasks required the processing of two stimuli throughout perception, memory and comparison stages of processing and so the set-size effect could be due to any one of these stages.

**Figure 12.**
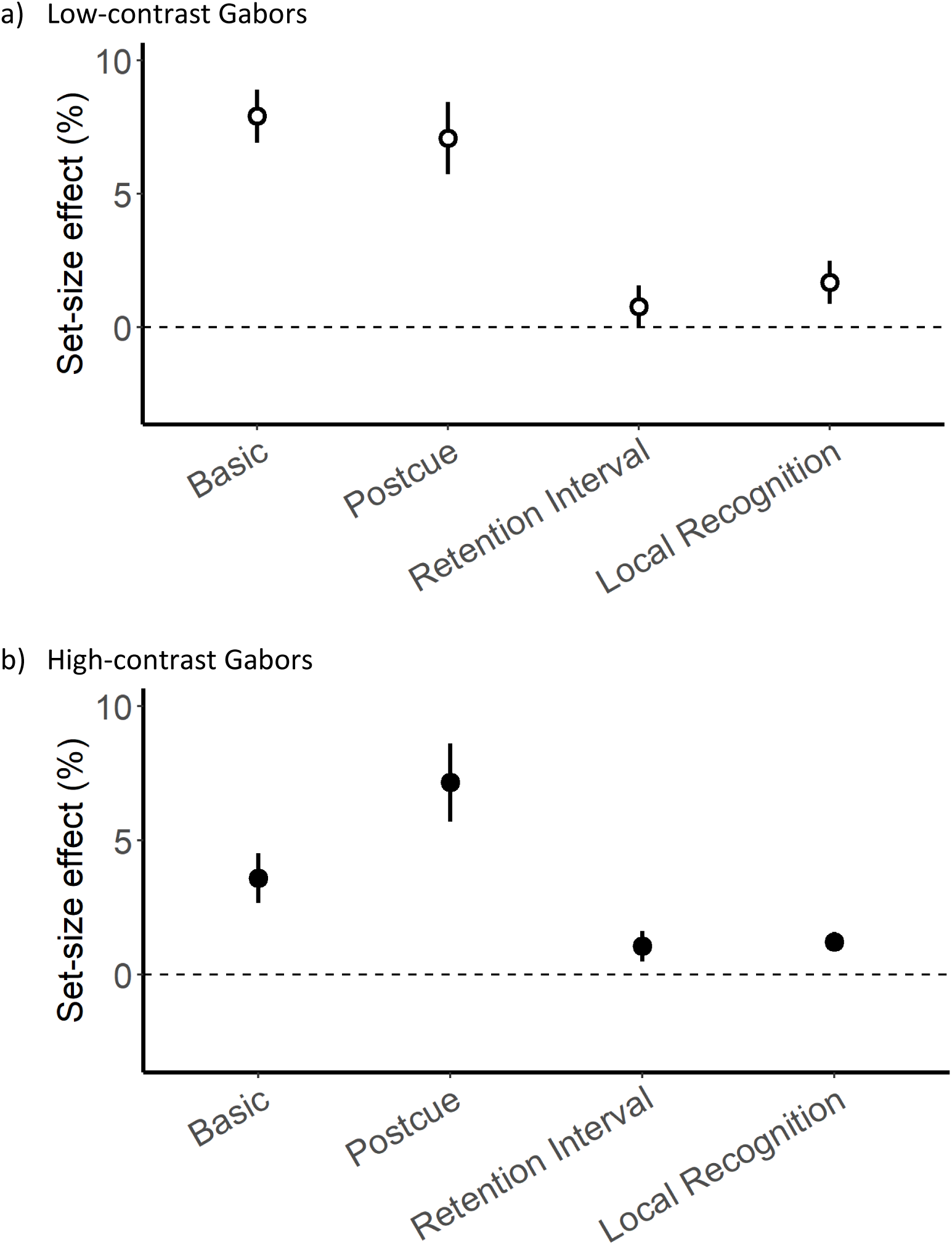
Magnitude of the effect of relevant set size for each of the experiments. The results for the low-contrast conditions are in Panel A and for the high-contrast conditions are in Panel B. Set-size effects are found for the basic and postcued experiments but little or no effect is found for the experiments using a retention-interval cue or local recognition.

For change detection using a retention-interval cue, there was little or no set-size effect: performance was the same, or nearly so, for two relevant stimuli as for one. For this task, two relevant stimuli must be processed by perception, memory encoding and satisfy the initial storage limits. But only a single relevant stimulus must be processed by memory storage processes (e.g. memory maintenance), memory retrieval and comparison.

For change detection using local recognition, there was little or no set-size effect. For this task, two relevant stimuli must be processed by perception, memory encoding and all aspects of memory storage. But only a single relevant stimulus must be processed by memory retrieval and comparison. Thus, these results are consistent with the capacity limit being due to memory retrieval or comparison.

The results for high-contrast stimuli typical of memory experiments are shown in Panel B. For both basic and postcued change detection there is a set-size effect. (There is a hint of a larger effect for postcued change detection compared to basic change detection, but further study is needed to confirm that detail.) For change detection using either a retention-interval cue or local recognition, there was little or no set size effect. Thus the pattern was similar for low and high-contrast stimuli.

### Implications for Perception

By perceptual processing, we mean the immediate processing of the stimulus rather than any delayed processing that is based on memory. It is challenging to separate effects of perception from effects of memory encoding and storage. In the experiments presented here it was unnecessary to pursue this distinction owing to the inference of unlimited capacity across all of these early processing stages.

Many change detection experiments vary the number of stimuli in the initial stimulus display and therefore change the sensory input from condition to condition which might introduce unintended effects, such as crowding (Parkes, Lund, Angelucci, Solomon, & Morgan, 2001). We avoided this potential confound by always having presenting the same stimuli in all conditions. Such constant displays rule out a nonattentional accounts such as crowding.

Finding no effect of relevant set size for perception of simple features is consistent with results from detection or detection-like tasks (Bonnel, Stein, & Bertucci, 1992; White, Runeson, Palmer, Ernst, & Boynton, 2017). These results differ from the predictions of theories that posit a limited capacity in perception for divided attention tasks. At one extreme are theories suggesting a bottleneck that allows only one stimulus to be identified at a time (Broadbent, 1958), while other theories are less severe suggesting a limited capacity or resource that is divided among relevant stimuli (e.g. Kahneman, 1973). These theories predict set-size effects because the stimuli are in competition for the limited processing capacity in perceptual stages. One explanation to these contrasting findings of limited and unlimited capacity for divided attention is a theory in which the outcome is dependent on the stimulus and task. Under some conditions, such theories predict unlimited capacity, and in other conditions they predict limited capacity (e.g. Hoffman, 1979; Scharff, Palmer, & Moore, 2011; Treisman & Gelade, 1980).

In summary, our results are inconsistent with a capacity limit in the perception of simple features. If the stimuli were not both perceived, then the experiments using a retention-interval cue or local recognition could not have improved performance and reduced the set-size effect. Thus, for the case of feature processing and 1 versus 2 stimuli, we can reject theories with limited-capacity perception (Kahneman, 1973; Pestilli et al., 2011). Of course, this does not rule out the possibility that these theories play a role with larger set sizes or for more complex stimuli.

### Implications for Perception of the Test Stimuli

There are two issues concerning the perception of the test stimuli in the second display. First, one might propose that limited capacity processing for the test stimulus contributes to the set-size effects. Particularly, because our experiments did not use long duration test displays. But, we argue that the case against capacity limits for the perception of the first stimulus display can be readily extended to the perception of the second display.

The second issue is that the perception the test display might interfere with memory of the first display. This possibility was suggested by Makovski & Jiang (2007). Recent studies have made a good case that such interference does contributed to the set-size effects and to the reduction of those set-size effects with the retention-interval cue (e.g. Souza, Rerko, & Oberauer, 2016). The idea that selective attention can protect against visual interference is similar to ideas on object substitution masking (Enns & Lollo, 1997).

For our study, there is no sign of this effect. It predicts an advantage for the condition with the retention-interval cue compared to local recognition. There was a similar lack of set-size effects for these two conditions. We suggest our study did not have this interference effect because our displays included dynamic visual noise for an extended time before and after both stimuli. This kind of filled interval is likely to cause its own visual interference (Makovski, Shim, & Jiang, 2006) and thus precluded additional interference from the test display. Thus, we argue that while interference from the test display undoubtable occurs in change detection, it had little role in the current study.

### Implications for Memory

In the general memory literature, the use of attentional cues to specify relevant and irrelevant stimuli (relevant set size) is best known as *directed forgetting*. For a review of this literature see MacLeod (1998). Most of this work has been for verbal stimuli, but see Williams, Hong, Kang, Carlisle, & Woodman (2013) and Hourihan, Ozubko, & MacLeod (2009). In this literature, there are three primary hypotheses to account for this kind of attentional effect in memory: selective rehearsal (Basden, Basden, & Gargano, 1993), retrieval inhibition (Bjork, 1989) or selective search in retrieval (Epstein, Massaro, & Wilder, 1972). Consider next specific memory hypotheses for the current study.

#### Memory encoding

Limited capacity at encoding means insufficient stimulus information is moved to storage. Sensory representations are retained only briefly after the stimulus event (e.g. Enns & Lollo, 1997; Palmer, 1988; Sperling, 1960) and therefore are unlikely to be encoded much beyond the first stimulus display in our task. The reduction of set-size effects by the retention-interval cue and by local recognition suggest that the stimulus representations are successfully encoded into memory storage because they are available for later access. This is consistent with those who have found insensitivity to stimulus array duration (Cowan et al., 2005; Luck & Vogel, 1997; Sperling, 1960), or have otherwise argued against capacity limits in encoding (Rideaux et al., 2015; Rideaux & Edwards, 2016). This result is not compatible with encoding that is serial or has limited capacity (e.g. Becker et al., 2013).

In summary, our results are inconsistent with a capacity limit in memory encoding. If the stimuli were not both encoded, then the retention-interval cue and local recognition could not have improved performance and reduced the set-size effect. Thus, for 1 versus 2 stimuli, we can reject theories with limited-capacity encoding (Becker et al., 2013; Palmer, 1990). Of course, this does not rule out the possibility that these theories play a role with larger set sizes or different encoding conditions.

#### Memory storage

There are several ways that memory storage might mediate set-size effects. The simplest is a limit on storage capacity itself. Such a limit might be in terms of the number of stimulus representations (e.g. Zhang & Luck, 2008) or in terms of the quality of the representations (e.g. Keshvari et al., 2013). These hypotheses are inconsistent with the results the experiments with a retention-interval cue or local recognition.

Two other ways that memory storage might mediate the set-size effect we have called selective maintenance and selective transfer. In selective maintenance, memory is improved by a limited-capacity maintenance process (e.g. rehearsal) that is applied to the representations of the relevant stimuli (e.g. Basden et al., 1993). In selective transfer, memory is improved by a limited-capacity process that transfers information about the relevant stimulus to a more durable memory storage (e.g. Sligte et al., 2008). Both of these hypotheses predict that retention-interval cues reduce set-size effects.

In a particularly relevant study by (Williams et al., 2013), one versus two colored squares were presented and the precision of recall was measured for a single color. Performance for a set size of two colors was worse than for one color, but was improved when a retention-interval cue indicated that only one color was relevant. The authors argued that results were evidence for a limited-capacity, selective maintenance process such as selective rehearsal.

The problem with selective maintenance and selective transfer is that neither predicts that set-size effects are reduced by using a local recognition task (Experiment 4). Thus, these hypotheses are inconsistent with our results. One possible reason for the lack of effects for selective transfer is that we used a largely noise-filled retention interval that might have eliminated contributions from less durable memory stores.

#### Memory retrieval

Limited capacity in retrieval means that although both stimuli were encoded and stored adequately, the representations from the first stimulus display are not retrieved with sufficient information for successful comparison (Shiffrin, 1970). Our results are consistent with a limit in retrieval. The reduction of the effect of relevant set size by a retention-interval cue or by local recognition allow one to make a single memory retrieval rather than two. If retrieval is imperfect or incomplete, it can account for both the set-size effect and its reduction with an appropriate cue.

Several hypotheses for incomplete retrieval have been suggested in the literature. By the *retrieval bottleneck hypothesis* (Carrier & Pashler, 1995; Oberauer, 2018), retrieval is a serial process: only one retrieval is possible at a time. For a brief test display, this can result an incomplete retrieval for the second stimulus to be retrieved.

By the *retrieval head start hypothesis* (Souza et al., 2016), retrieval requires time so that a retention-interval cue provides a head start. This hypothesis has also been supported by the finding that delaying the response improves performance.

By the *long-term-memory retrieval hypothesis* (Beck & van Lamsweerde, 2011), explicit retrieval cues such as the retention-interval cue or the single test stimulus in local recognition, improve retrieval for long-term memory that supplements retrieval based on working memory alone.

By the *retrieval inhibition hypothesis* (Bjork, 1989), the cues improve retrieval by inhibiting the memory of irrelevant stimuli.

By the *selective retrieval hypothesis* (Epstein et al., 1972), the cues provide additional context that helps guide the retrieval of the relevant memory. One interesting variation on this idea is that the more specific context protects the retrieval process from interference from the other stimuli (Oberauer & Lin, 2016). In summary, there is no shortage of hypotheses about how retrieval might modulate set-size effects in change detection.

These retrieval accounts clash with accounts of working memory that assert no retrieval for the stimulus representations held in the *focus of attention*. By this account, memories for the stimuli in the focus of attention can be directly accessed for comparison to the stimuli in the second display. McElree (1998) has argued that this focus of attention is limited to as little as one object while Cowan (1988) has argued that it can encompass several objects. In more recent reviews, this proposal has been defended by Cowan (2011) and has been criticized by Oberauer (2013) and the debate continues (e.g. Vergauwe & Langerock, 2017). If direct access to multiple objects is found to hold, then retrieval cannot be the limit for the results found here.

### Implications for Comparison

The results with the retention-interval cue and with local recognition are also consistent with a limit in the comparison process. Only one comparison has to be made for these conditions. This hypothesis is supported by results showing that despite failing to detect a change, a subsequent probe about stimulus identity demonstrates sufficient information was available in memory (Angelone et al., 2003; Farell, 1985; Fernandez-Duque & Thornton, 2000; Hyun et al., 2009; Mitroff et al., 2004; Simons et al., 2002). For example, Mitroff et al. (2004) showed that, despite encoding sufficient information about all of the relevant stimuli for a 2AFC task asking whether stimuli had been present in either display, observers still failed to detect changes. Findings such as this have been interpreted as that one can retrieve the relevant memory, but fail to make the correct comparison when there are multiple comparisons to be made.

### Implications for Decision

Decision can contribute to set-size effects when there is uncertainty in mapping stimuli to a specific response. As the number of relevant stimuli increases, additional uncertainty from each stimulus is included in the decision which limits performance (Palmer et al., 2000; Sperling & Dosher, 1986; Swets et al., 1961). Change detection tasks are often structured such that all stimuli contain relevant information that must be integrated to make the decision. In Experiment 1, for example, observers were asked to detect a change occurring anywhere in the array, so that all locations are informative to the decision. In Experiment 2, by contrast, the postcue allowed observers to map a single stimulus to their judgement.

In fact, the addition of a postcue did not reduce set-size effects. Similar results have been found in previous change detection experiments with colored squares (Luck & Vogel, 1997; Wheeler & Treisman, 2002) and with letters (Becker, Pashler, & Anstis, 2000). In contrast to these results, there is evidence of postcue effects in a study of Gabor patches (Wright, Green, & Baker, 2000). So the results with simple stimuli are unclear.

A somewhat different pattern of results have been found for two studies using familiar objects. Postcues reduced the set-size effects in an experiment using an array of familiar objects (Beck & van Lamsweerde, 2011) and for an experiment using familiar objects in natural scenes (Hollingworth, 2003). These experiments also used relatively long study display durations to encourage the use of long-term memory. In addition, Beck and van Lamsweerde provide specific evidence for the role of long-term memory and argue that the postcue effects are due to encouraging retrieval from long-term memory.

There is an alternative view of the comparison process worth mentioning. In Experiment 2, a correct response requires accurate information about the location of the change as well as whether a change occurred. If location information was imperfect, one would expect a decline in performance in Experiment 2 compared to Experiment 1. That was not found. The analysis of congruency effects is relevant to this possibility. If location information is unreliable, then one would expect better performance incongruent trials than in incongruent trials. For congruent trials, both locations require the same response so unreliable information about location does not affect performance. That is not true for incongruent trials. In fact, there were no congruency effects in any of the experiments. This is consistent with location information not limiting performance in these experiments. This is perhaps not surprising since the differences in location were maximized (left versus right side of fixation) and the use of relevant set size minimizes changes in context.

Why might there be no effect of set size on decision in the current experiments? One possibility is that these memory tasks depend on discrete representations rather than the continuous representations as assumed by typical signal detection models. A simple two-state representation does not predict an uncertainty effect on decision (see the high threshold theories described in Palmer et al., 2000). This idea is supported by a recent study of color change detection task by Rouder et al. (2008). See also related studies of long-term memory (Province & Rouder, 2012). Unfortunately, the evidence for this possibility is not universal. Ricker, Thiele, Swagman, & Rouder (2017) used similar methods on an orientation-based change detection task and found no evidence for a discrete representation. Thus, the discrete representation model appeared to be a viable account of the results found here for decision, but there is doubt that they are general to all change detection tasks.

### Generalization

In this article, we focus entirely on the case of orientation, but all of our interpretation has assumed that this is not a special feature. Many studies of change detection use salient and highly discriminable colors as the feature and thus color would therefore be a natural direction for generalization. In favor of similarity between our stimuli and highly discriminable colors, there were the same pattern of results in our high-contrast condition as our low-contrast condition. This is consistent with our results not being due to limited discriminability. The question of whether color is different from orientation is less clear. In a series of papers asking whether encoding into visual short-term memory occurs in series or in parallel, Becker and colleagues found color and orientation to be different (Becker et al., 2013; Liu & Becker, 2013; Mance, Becker, & Liu, 2012; Miller, Becker, & Liu, 2014). In a local recognition task, they compared performance for stimuli presented simultaneously versus sequentially. If encoding has unlimited capacity then there should be no difference between these conditions, but if encoding has limited capacity then performance should be better in the sequential condition. They found that for color – but not orientation – the performance was equivalent between the simultaneous and sequential conditions. They suggested that color, but not orientation, can be encoded in parallel for one versus two stimuli. What to make of these results? First, these results suggest that orientation and color are sometimes processed differently. Second, the sequential advantage for orientation is inconsistent with our data. In a local recognition task, there was unlimited capacity for two stimuli. The differences found by Becker and colleagues are in the wrong direction to predict a different pattern of results for color in our task given that they find that encoding color has unlimited capacity. Thus, we think it likely that the results found here for orientation will generalize to color.

Our use of a coarse change in orientation of 90° means that there can be considerable loss in the precision of memory without any impact on performance accuracy. With a finer discrimination task, one might find that perception is not unlimited, and that precision decreases with one to two stimuli as others have found (Keshvari, van den Berg, & Ma, 2012). On the other hand, there are other results that are consistent with unlimited capacity for precise feature judgments (e.g. Scharff et al., 2011). Thus, this issue is yet to be resolved.

An incidental feature of our experiments was the use of *1/f* noise during much of the retention interval. Such pattern noise might have minimized the role of less durable memory stores. It is possible that the use of filled intervals will prove useful to simplify the memories underlying change detection.

Many visual memory studies focus on limits occurring at larger set sizes, such as two versus four stimuli. It is possible that additional limits are introduced with more stimuli. For example, crowding becomes an expected perceptual limit with more than a few stimuli (e.g. Parkes et al., 2001), and memory storage for more than three or four objects also has been discussed extensively as a limit (e.g. Cowan, 2000; Landman et al., 2003; Luck & Vogel, 1997). It remains to be seen whether our account of set sizes 1 and 2 generalizes to larger set sizes.

## Conclusion

We investigated set-size effects in change detection. Our goal was to find the locus of the initial capacity limits revealed by set sizes 1 and 2. Relevant set size was used rather than display set size to measure purely attentional effects and not other phenomena such as crowding. For basic change detection, there was an effect of relevant set size: performance was worse with two relevant stimuli compared to a single relevant stimuli. This effect was also found for postcued change detection. But the results were different for an experiment with a retention-interval cue between the stimulus displays, and for an experiment that used local recognition to test memory for a single stimulus. For these two experiments, there was little or no set-size effect. We infer that the capacity limit with just two stimuli is due to memory retrieval and/or comparison. For these set sizes, perception, memory encoding and storage are ruled out as the locus for the capacity limit. This result for 1 versus 2 stimuli, is inconsistent with a variety of theories including limited-capacity perception for all stimuli (e.g. Kahneman, 1973; Pestilli et al., 2011) and limited-capacity memory storage (e.g. Keshvari et al., 2013; Zhang & Luck, 2008).

## Author Note

We thank Cathleen M. Moore and Alex L. White for insightful suggestions on these studies. This research was partly supported by a grant EY12925 to G. M. Boynton from the National Institute of Health.

## Open Practices

Data from this study will be available in the repository of the Open Science Framework at: (under construction as of January 2020). Other materials are available on request.

## Appendix: Further analyses of all experiments

### Stimulus timing

For all experiments other than Experiment 3, the second stimulus and postcue occurs beyond the time limits of iconic memory (Sperling, 1960). In Experiment 3, however, the retention-interval cue appears 250 ms after the end of the stimulus display. The Gabor could be present up to the last 200 ms of the stimulus display, resulting in the possibility that in some trials the retention-interval cue could appear within 450ms. We suspect this is beyond the limits of iconic memory for our displays, but what if it isn’t? It would then be possible for the observer to encode only one stimulus from the first display and lead to improved performance in the cue-both condition relative to other experiments.

To test the possibility that stimulus timing affected performance we correlated the timing of the stimulus in the first stimulus display with the performance in that trial. If the presence of the stimulus in iconic memory at the time of the retention-interval cue was of benefit then performance should improve the later in the trial the Gabor was presented (i.e. a positive correlation). None of our observers showed a significant correlation in Experiment 3 or any other experiment (mean correlation coefficient = .01, *p* > .1 with Bonferroni correction). Thus, there is no evidence of an effect of when the retention-interval cue appeared relative to the stimuli.

### Congruency

The results of dual task experiments (2, 3, and 4) can be further broken down by congruency of the stimuli events at each location on each trial. Congruent trials are ones in which the same stimuli event (e.g. a change) occurred at both locations. Incongruent trials are ones in which different stimuli events happened at each location (e.g. a change on one side, and not on the other). The presence or absence of congruency effects indicates the degree of independence across locations. If performance is independent then there should be no difference in performance between congruent and incongruent trials. On the other hand, if there is a trial-by-trial dependency across sides then there should be a difference in performance between congruent and incongruent trials (e.g. Bonnel, Stein, & Bertucci, 1992). Such an effect is indicative of a divided attention effect beyond an accuracy difference across set-size conditions. Examples of congruency effects have been found in some but not all divided attention experiments (e.g. Navon & Miller, 1987). The congruency effects for the three dual task experiments (Experiments 2-4) are shown in Figure 13. There was no reliable effect of congruency in any experiment or contrast condition.

**Figure 13.**
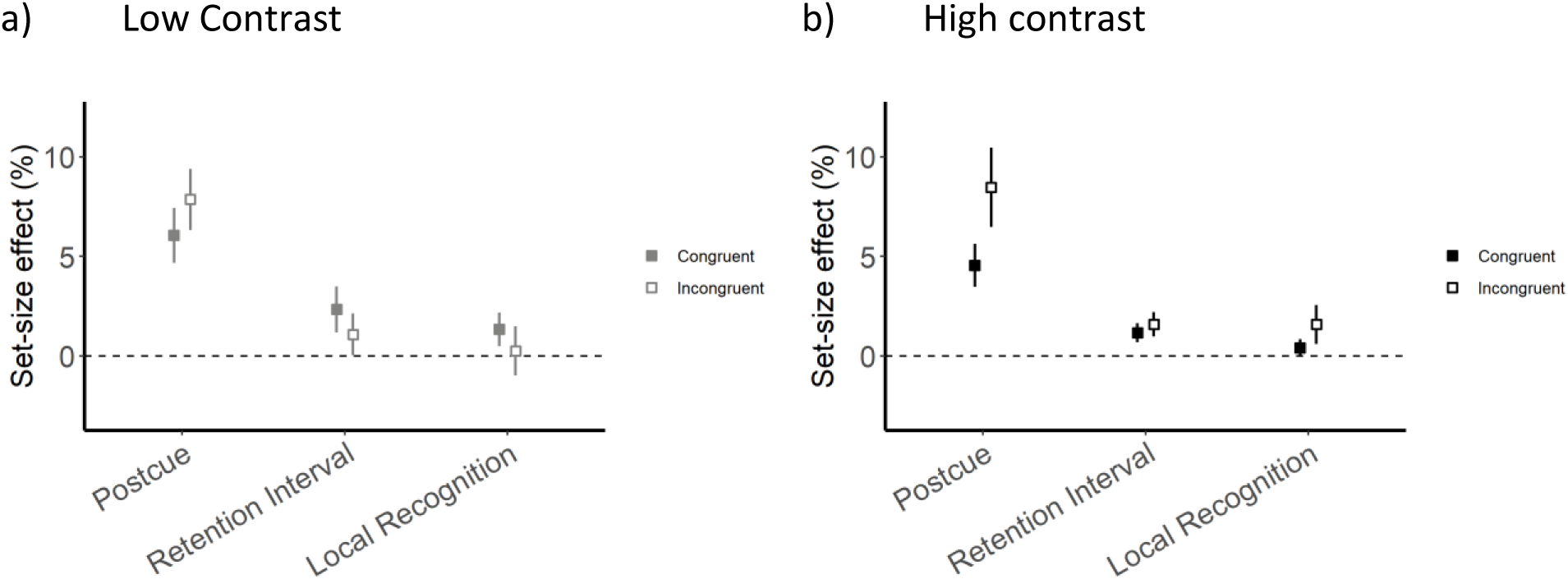
Congruency results for postcued change detection, retention-interval cued change detection and local recognition at (a) low and (b) high contrast. There was no sign of congruency effects in these experiments.

### Orientation similarity and perceptual grouping

There are several findings that suggest perceptual grouping did not play a significant role in the present experiments. First, performance was no better or worse in local recognition than with the retention-interval cue. Removing the reference of the second side stimulus in local recognition should have removed any benefits of perceptual grouping because the context has changed. This predicts a smaller set-size effect in retention-interval cued change detection. However, the set-size effect was similar in these two experiments suggesting that perceptual grouping was not improving performance for the set size 2 conditions with the retention-interval cues relative to local recognition.

Second, if certain sets of orientations were more conducive to grouping then there should be differences between these pairs. There were two possible differences between angles on the two sides (22.5° and 67.5°) and the mean performance was the same in each case (22.5°: mean = 86.1, se = 0.4; 67.5°: mean = 86.1, se = 0.4). A three-way ANOVA 4 (experiment) x 2 (contrast) x 2 (orientation distance) found main effects of experiment and contrast but not of orientation difference, F(1, 179) = 0.71, *p* = .40.

Third, when the side of the change is not relevant, as in Experiment 1, perceptual grouping might improve performance if any stage of perception, memory or comparison was limited because it is only the one grouped set of stimuli that must be encoded, stored and retrieved for comparison. Any change in this stimulus from the first to the second display would be reported as a change. By making the locations independent in Experiment 2 this strategy is less helpful. The magnitude of the set-size effect was similar in Experiment 1 and 2 suggesting that perceptual grouping was not helpful as a strategy for reducing the effective number of stimuli.

Why are there no effects of perceptual grouping? Any potential benefit of perceptual grouping might have been eliminated due to the presence of noise or the randomized location of the stimuli from first to second stimulus display.

